# Morphine-context associative memory and locomotor sensitization in mice are modulated by sex and context in a dose-dependent manner

**DOI:** 10.1101/2023.11.03.565492

**Authors:** Peter U. Hámor, Matthew C. Hartmann, Aaron Garcia, Dezhi Liu, Kristen E. Pleil

**Author notes:** Declaration of interests: The authors declare that they have no known competing financial interests or personal relationships that could have appeared to influence the work reported in this paper.

## Abstract

Sex differences in opioid use, development of opioid used disorder, and relapse behaviors indicate potential variations in opioid effects between men and women. The locomotor and interoceptive effects of opioids play essential roles in opioid addiction, and uncovering the neural mechanisms underlying these effects remain crucial for developing effective treatments. In this study, we examined the dose-dependent effects of morphine on locomotor sensitization and the strength and stability of morphine-context associations in the conditioned place preference (CPP) paradigm in male and female mice, as well as the relationships between these measures. We observed that while CPP is similar between sexes, the locomotor effects of repeated morphine administration and withdrawal differentially contributed to the strength and stability of morphine-context associations. Specifically, females exhibited higher morphine-induced hyperlocomotion than males regardless of the context in which morphine was experienced. Greater locomotor sensitization to morphine in females than males emerged in a dose-dependent manner only when there was sufficient context information for CPP to be established. Additionally, the relationships between the locomotor effects of morphine and the strength and stability of CPP were different in males and females. In females, positive acute and sensitizing locomotor effects of morphine were correlated with a higher CPP score, while the opposite direction of this relationship was found in males. These results suggest that different aspects of the subjective experience of morphine intoxication and withdrawal are important for morphine abuse-related behaviors and highlight the importance of sex-specific responses in the context of opioid addiction.

## 1 Introduction

More than 75,000 people in the United States die every year from opioid use and overdose, and this trend continues to rise (Spencer *et al*., 2022). These deaths are associated with an increase in opioid use disorder (OUD), a chronic, relapsing disorder characterized by compulsive drug seeking and dysregulation of brain circuits underlying motivational states, reward, and drug withdrawal-induced negative affective states (Koob & Volkow, 2016). Sex differences in opioid use, the development of OUD, and the drivers of relapse behavior suggest that the pharmacological actions of opioids may differ between men and women across stages of opioid use (Fillingim *et al*., 2003; Back *et al*., 2010; Jamison *et al*., 2010; Darnall & Stacey, 2012; Manubay *et al*., 2015; Fullerton *et al*., 2018; Marsh *et al*., 2018; Silver & Hur, 2019). Studies show a more rapid progression from initial opioid use to opioid abuse in women than men (called telescoping of opioid use), even though the overall prevalence of substance use disorder (SUD; nida.nih.gov, 2020), as well as OUD, is higher in men (Silver & Hur, 2020). There are also many other sex differences in relapse-related behaviors, including craving in response to different types of drug-related cues (Yu *et al*., 2007), however, behavioral and subjectively evaluated results of human studies often contradict each other. Nonetheless, there is growing recognition of sex differences in the behavior and biology of opioid exposure and use that are important for addressing more effective treatment approaches (Lee & Ho, 2013; Becker & Chartoff, 2019). Recent preclinical evidence also demonstrates sex differences in the nuances of opioid addiction (Bravo *et al*., 2020; Zhang & Kreek, 2020), including the observation that female mice are more likely to use opioids, such as fentanyl, regardless of the negative consequences of their use (Monroe & Radke, 2021), a behavior associated with OUD in humans (Guha, 2014). While these studies suggest differences between males and females in the effects of opioids that could have significant consequences, little else has been examined in females.

The locomotor and interoceptive effects of opioids are two distinct but interconnected phenomena that play a critical role in the development of opioid addiction. Locomotor sensitization, characterized by an increase in locomotor activity following repeated exposure to opioids, has been shown to be related to some drug-seeking and drug-taking behaviors for several classes of drugs, including opioids (Robinson & Berridge, 2000). Interoceptive effects, such as the subjective sensations produced by opioids such as euphoria, are more directly linked to drug reward and play a crucial role in the development of opioid addiction (Koob & Volkow, 2010). Both the locomotor and interoceptive/reinforcing effects of opioids are heavily modulated by common neuromodulators and pathways, such as the midbrain/mesolimbic dopaminergic system (Wise, 2008). Dopamine release is associated with the subjective experience of opioid reward and the locomotor effects of opioids, depending on the subregion and projection target(s) of the originating midbrain dopamine neurons (Robinson & Berridge, 2000; Koob & Volkow, 2010). While both contribute to drug seeking and taking behaviors, the locomotor effects of opioids do not always correspond to the subjective experience of drug exposure, suggesting they cannot be used as the singular proxy for opioids’ rewarding effects (Bardo & Bevins, 2000). Understanding the relationship between these two phenomena is critical for elucidating the neural mechanisms underlying opioid addiction and developing effective treatments for this disorder.

These effects of opioids may play important roles in the formation of learned associations between opioids and the context in which they are experienced, a critical component of the development and maintenance of OUD (Robinson & Berridge, 1993; Koob & Volkow, 2010). The conditioned place preference (CPP) paradigm provides a useful framework for understanding the relationships between the interoceptive and locomotor effects of drugs and withdrawal from them. In the paradigm, drug exposure is paired with a specific context whereas the lack of drug (control vehicle) is paired with a different specific context, within the same broader environment, resulting in a conditioned preference for the drug-paired context on a subsequent session in which no drug or vehicle is administered (Tzschentke, 2007). While drug-context associations have been thought to be primarily driven by the rewarding effects of the opioid drug (Mucha *et al*., 1982), the context associated with opioid withdrawal-related aversion may also be important for driving preference for opioid-paired context(s) (Koob & Le Moal, 2001). Females display increased morphine/opioid-induced hyperlocomotion and more severe physical signs of protracted withdrawal than males (Bravo *et al*., 2021), which may contribute to sexually dimorphic behavioral mechanisms underlying the strength of opioid-context associations. Here, we examine the dose-dependent effects of morphine on locomotor sensitization and the strength and stability of morphine-context associations in the CPP paradigm in male and female mice, as well as the relationships between these measures.

## 2 Material and methods

### 2.1 Animals

Female and male C57BL/6J mice were purchased from The Jackson Laboratory (Bar Harbor, ME, USA) and group housed in standard cages with 5 mice/cage with *ad libitum* access to food and water in a colony room on a 12:12 hr light-dark cycle. Experiments began approximately 3h into the dark phase of the cycle. All experimental procedures were approved by the Institutional Animal Care and Use Committees at Weill Cornell Medicine.

### 2.2 Conditioned place preference

The CPP apparatus consisted of a standard mouse arena (27.3×27.3×20.3 cm) separated into two compartments by a two chamber place preference insert with removable guillotine door (Med Associates Inc., Fairfax, VT, USA). The left compartment had white walls with horizontal black stripes, and the right compartment had white walls with vertical zig zag black lines. In the low context version of the assay, both compartments had white, smooth plexiglass flooring. In the high context version of the assay, the right compartment remained the same, but the left compartment had textured flooring to add a distinct contextual feature (**Fig. 1B**). Mice were habituated to handling by the experimenter for five days prior to the experiment. Each day of the paradigm, mice were allowed to habituate to the testing room for 60 min prior each testing or conditioning session. On Day 1, mice underwent a pretest in which they were placed in the apparatus and allowed to freely explore for 15 min, with no injections administered prior to the test. No mice in this study displayed an initial compartment preference of greater than 70% that would have been considered a bias requiring exclusion (Goltseker & Barak, 2018). The following day, mice began the four-day conditioning phase of the experiment. On each of conditioning days 1-4 (C1-C4), mice underwent a morning session in which they were injected with 0.9% sterile saline (*i.p.*) and confined to one of the two compartments of the apparatus for 30 min, pseudo-randomly assigned regardless of initial preference; six hrs later, mice underwent an afternoon session in which they were injected with morphine sulfate (5, 10, 20, or 50 mg/kg in 0.9% sterile saline) and confined to the other compartment of the apparatus. The next day (“post-test day 1”, or P1), mice underwent another test session identical to the pretest. Mice then remained in their home cages except to be tested approximately every 10 days through P70, and then again at P100. The animal’s position was tracked via a 16×16 grid of infrared beams and Med Associates software calculated locomotion and time spent in each compartment. CPP score was calculated as the difference between the time spent in the morphine-paired compartment of the apparatus on each post-test minus that on the pretest.

**Figure 1:**
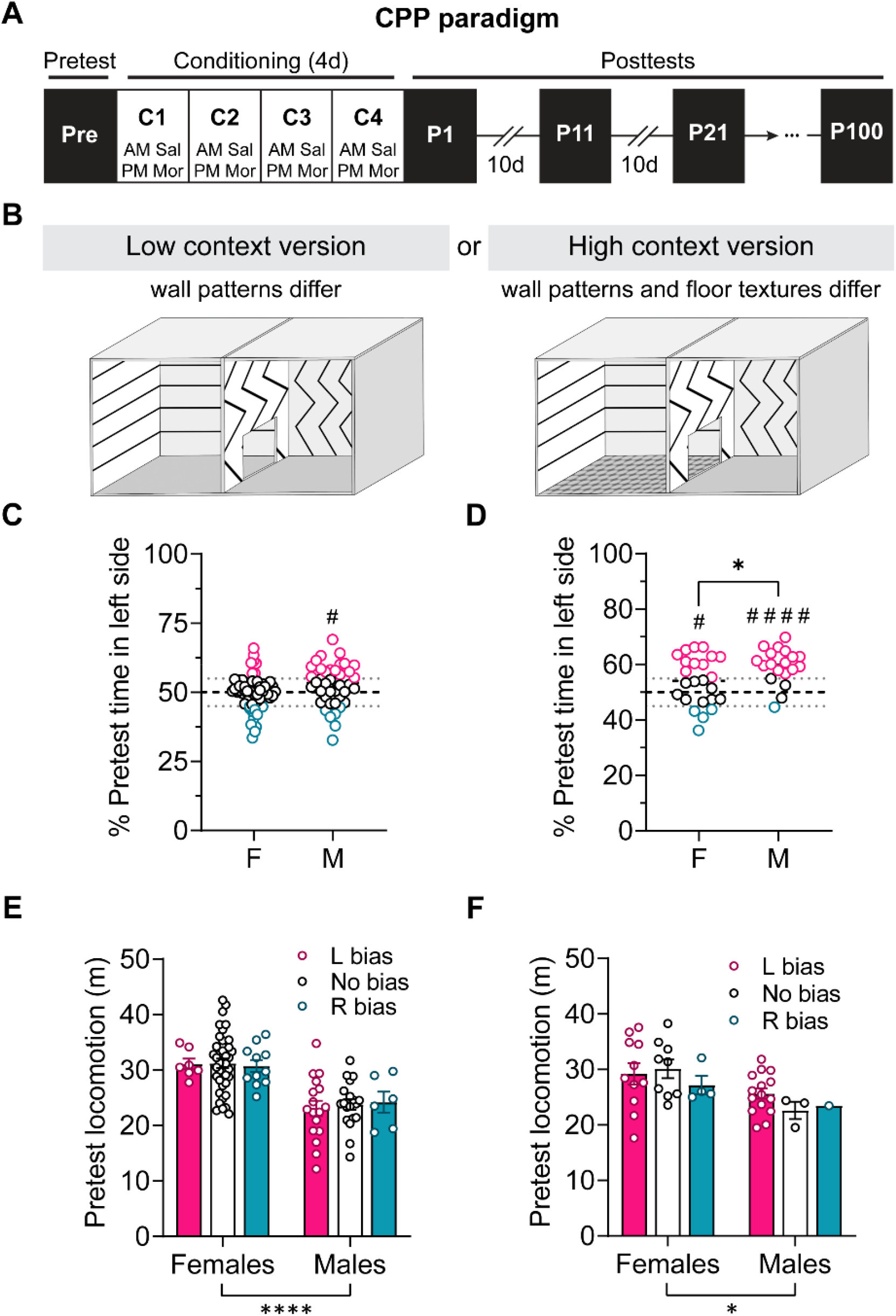
Sex-dependent preference for context elements. **A)** Experimental timeline of the conditioned place preference (CPP) paradigm. **B)** Illustration of the CPP apparatus in the low and high context version of the apparatus. In the low context version, the walls were distinctly patterned with horizontal stripes in the “left” compartment and zig zag stripes in the “right” compartment, but the flooring was smooth in both compartments. In the high context version, the flooring was also distinct between compartments, with the “left” compartment altered to a textured rubber flooring to distinguish it tactilely from the “right” compartment smooth flooring. **C-D)** % Time spent in the left compartment of the apparatus in the low (**C**) and high (**D**) context versions of the apparatus on the pretest, when mice could freely explore both compartments of the apparatus for the first time. Individual data points are color coded to reflect categorization of compartment bias on the pretest (pink: > 55%; teal: < 45%; black: between 45% and 55% in left compartment). *p < 0.05 for unpaired t-tests between males and females; ^#^p < 0.05, ^####^p < 0.0001 for one-sample t-tests within group compared to the null hypothesis mean of 50%. **E-F)** Pretest locomotion in females and males, subcategorized by compartment bias of 55% as defined in **C and D**. Females had greater pretest locomotion than males in both the low (**E**) and high (**F**) context versions of the assay, but locomotion was unaffected by compartment bias. *p < 0.05, ****p < 0.0001 for main effects of sex in 2×2 ANOVAs.

### 2.3 Statistical analysis

Statistical analyses were done using GraphPad Prism 9 (GraphPad Software, San Diego, CA, USA) with an alpha level of 0.05. Gaussian distributions within and equality of variance between groups were confirmed, zero outliers detected and excluded using Q-Q plots, and parametric statistics were performed on all data. Analysis of variance (ANOVA) and repeated measures ANOVA (RM-ANOVA) were used to evaluate the effects of sex, compartment, morphine dose, test day, and other variables on locomotion and CPP. Significant effects and interactions were followed up with unpaired or paired t-tests with Holm-Šidák corrections for multiple comparisons, and multiplicity-adjusted p values are reported. One-sample t-tests were used to evaluate whether CPP scores were significantly different from chance. A total of 12 mice (5 females, 7 males) were excluded from the analysis due to arena IR beam malfunctioning that prevented acquisition of accurate tracking data. No other animals were excluded for any other reason.

## 3 Results

### 3.1 Sex-dependent preference for context elements

Female and male mice were tested in two versions of a two-compartment conditioned place preference (CPP) assay (**Fig. 1A,B**). We assessed whether male and female mice displayed innate preferences for context features during their first exposure to the CPP apparatus on the pretest, and further whether these preferences affected their locomotor behavior (**Fig. 1C-F**). In the low context assay, in which the two compartments only differed in the pattern of stripes on the walls, females displayed no initial compartment bias on the pretest, with a mean of 50.0% time spent in each compartment (**Fig. 1C**; one-sample t-test: t(57) = 0.01, p = 0.996). Males spent more time in the “left” compartment with horizontal stripes (52.7%; one-sample t-test: t(41) = 2.37, ^#^p = 0.022), however this was not statistically different from females (t(98) = 1.97, p = 0.051). In the high context version of the assay in which both the wall pattern and floor texture differed, both sexes spent more time in the “left” compartment with horizontal wall stripes and textured flooring (compared to zig zag wall and smooth flooring; one-sample t-tests: F: t(23) = 2.27, ^#^p = 0.033; M: t(18) = 6.60, ^####^p < 0.0001; **Fig. 1D**), and this compartment preference was higher in males vs. females (59.5% vs. 54.0%; t(41) = 2.31, *p = 0.026). Nonetheless, no mice spent more than 70% of their time in either compartment, a common exclusion criterion used for CPP studies (Goltseker & Barak, 2018). In addition, a preference for the left or right compartment during the pretest, defined by >55% time spent in that compartment, did not affect locomotion in either sex on either version of the assay (**Fig. 1E-F**). In the low context assay, a 2×2 ANOVA showed a main effect of sex (F (1, 93) = 38.25; ****p < 0.0001) but not compartment bias (F (2, 93) = 0.05200, p = 0.949) and no interaction between the two (F (2, 93) = 0.1063, p = 0.899) on pretest locomotion (**Fig. 1E**). A similar result emerged in a 2×2 ANOVA for the high context version of the assay, with an effect of sex (F (1, 37) = 5.146, *p = 0.029) but not compartment bias (F (2, 37) = 0.3800, p = 0.687) or interaction (F (2, 37) = 0.5708, p = 0.570) on pretest locomotion (**Fig. 1F**). These results show that females have higher locomotion in novel contexts, as previously described in the literature (Borbélyová *et al*., 2019), but that small sex-dependent biases for context elements in the CPP task do not alter locomotion in these contexts, *per se*.

### 3.2 Effects of sex and context on the acute locomotor effects of morphine

We next examined the acute effects of morphine on locomotion in the low and high context versions of the CPP assay. We evaluated whether the concentration of morphine and the compartment of the CPP apparatus in which mice were confined following morphine or saline on conditioning day 1 (C1), which was randomly assigned, affected locomotion differently in males and females. In both sexes and in both contexts, we found a dose-dependent increase in morphine-induced hyperlocomotion that did not depend on the compartment in which morphine was experienced for the first time on C1; we found no effects on locomotion following saline injections (Fig. 2). In the low context version of the CPP assay in females (Fig. 2A), a 2×2 repeated measures (RM) ANOVA revealed a main effect of morphine dose (F (3, 50) = 10.74, ****p < 0.0001) but no effect of apparatus compartment (F (1, 50) = 0.04, p = 0.834) or interaction (F (3, 50) = 0.26, p = 0.851) on morphine-induced hyperlocomotion. *Post hoc* comparisons with H-S corrections between morphine doses showed that 20 mg/kg and 50 mg/kg locomotion were significantly higher, and 10 mg/kg trended toward being higher, than 5 mg/kg (10: t(50) = 2.54, ^%^p = 0.056; 20: t(50) = 5.11, ^$$$$^p < 0.0001; 50: t(50) = 4.35, ^$$$^p = 0.0003). In addition, 20 mg/kg and 50 mg/kg locomotion trended toward being higher than 10 mg/kg (20: t(50) = 2.27, ^&^p = 0.056; 50: t(50) = 2.46, ^&^p = 0.056) but did not differ from each other (t(50) = 1.14, p = 0.262). A 2×2 RM-ANOVA showed that post-saline locomotion on C1 did not differ between compartments or morphine dose (ps > 0.15). A similar pattern emerged in the high context version of the assay in females (Fig. 2B). A 2×2 RM-ANOVA revealed a main effect of morphine dose (F (1, 19) = 8.37, **p = 0.009) but no effect of compartment (F (1, 19) = 0.13, p = 0.722) or interaction (F (1, 19) = 0.004, p = 0.950) on morphine-induced hyperlocomotion. A *post hoc* comparison between 10 and 20 mg/kg, regardless of conditioning compartment, showed that morphine-induced hyperlocomotion was greater for 20 mg/kg than for 10 mg/kg (t(21) = 3.11, ^##^p = 0.005). Again, locomotion on C1 following saline injection was unaffected by compartment or morphine dose (ps > 0.05). In males, we also observed a dose-dependent effect of morphine, with no effect of conditioned compartment, on locomotion. In the low context version of the assay (Fig. 2C), a 2×2 RM-ANOVA revealed a main effect of morphine dose (F (3, 39) = 4.88, **p = 0.006) but no effect of compartment (F (1, 39) = 0.004, p = 0.949) or interaction (F (3, 39) = 0.80, p = 0.499) on morphine-induced hyperlocomotion. *Post hoc* comparisons with H-S corrections between morphine doses showed that 20 mg/kg and 50 mg/kg locomotion were significantly higher than 5 mg/kg (20: t(39) = 3.37, ^$^p = 0.010; 50: t(39) = 2.75, ^$^p = 0.044; all other ps > 0.10). A 2×2 RM-ANOVA showed that post-saline locomotion on C1 was unaffected by compartment or morphine dose (ps > 0.05).

**Figure 2:**
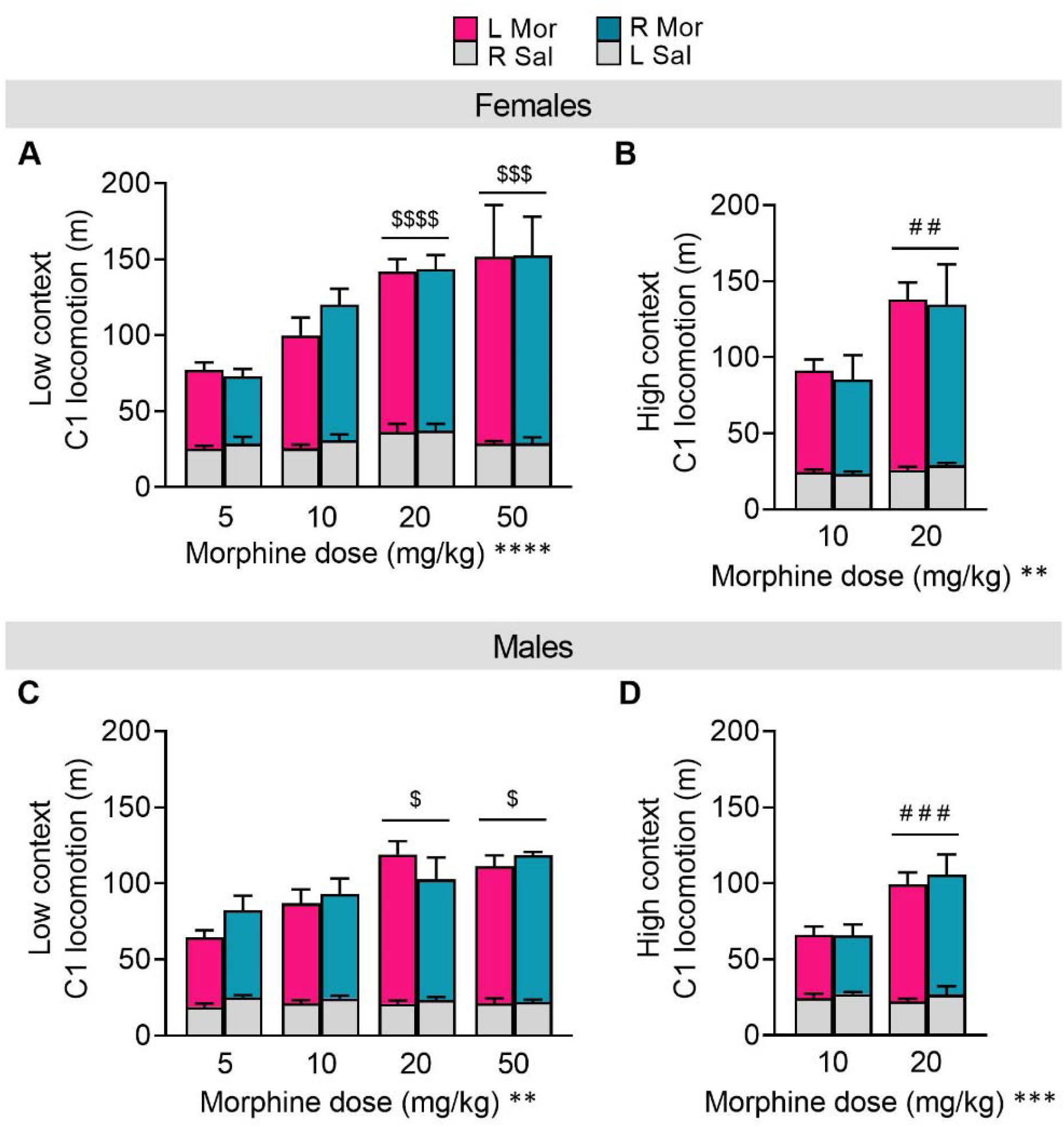
Dose-dependent acute locomotor sensitivity to morphine is unaffected by the context with which morphine is paired. **A-B)** On the first day of conditioning (C1) in the low context (**A**) and high context (**B**) versions of the CPP assay, females displayed acute hyperlocomotion following the morphine injection on the afternoon conditioning session, regardless of whether the morphine was paired with the left or right compartment; this locomotor sensitivity was dose-dependent, as cohorts receiving higher doses of morphine displayed greater morphine-induced hyperlocomotion. Locomotion following saline injection on the morning session (6 hr earlier) was similar across morphine dose groups and paired compartment. **C-D)** Males displayed a similar dose-dependent hyperlocomotor response to morphine that was similar in both compartments of the apparatus in the low (**C**) and high (**D**) context assays, with no differences in saline locomotion. **p < 0.01, ****p < 0.0001 indicate main effect of morphine dose in 2×2 RM-ANOVAs; ^$$$^p ≤ 0.001, ^$$$$^p < 0.0001 indicate difference from 5 mg/kg morphine dose, ^##^p ≤ 0.01, ^###^p ≤ 0.001 indicates difference from 10 mg/kg morphine dose in *post hoc* t-tests between doses.

Similarly, in the high context assay (Fig. 2D), a 2×2 RM-ANOVA revealed a main effect of morphine dose (F (1, 16) = 19.76, ***p = 0.0004) but no effect of compartment (F (1, 16) = 0.006, p = 0.939) or interaction (F (1, 16) = 0.09, p = 0.771) on morphine-induced hyperlocomotion. A *post hoc* comparison between 10 and 20 mg/kg, regardless of conditioning compartment, showed that morphine-induced hyperlocomotion was greater for 20 mg/kg than for 10 mg/kg (t(18) = 4.73, ^###^p = 0.0002). Again, locomotion on C1 following saline injection was unaffected by compartment or morphine dose (ps > 0.20). These results show that males and females display similar dose-dependent morphine-induced hyperlocomotion that is not affected by context elements present in the apparatus when mice first experience morphine. Together with results from Fig. 1, we also find that both basal and saline-induced locomotion do not depend on the innately preferred compartment of the apparatus, nor the compartment paired with morphine/saline.

### 3.3 Effects of context on sex differences in morphine-induced locomotor sensitization

Following the assessment of the acute locomotor effects of morphine, we examined locomotor sensitization to morphine across daily conditioning sessions (Fig. 3). We first measured locomotion during morning saline sessions across conditioning days in the low context version of the assay (Fig. 3A). For saline conditioning session locomotion in the 5 mg/kg morphine dose, a 2×2 RM-ANOVA showed a main effect of sex (F (1, 18) = 5.37, *p = 0.033) and a trend for effect of conditioning day (F (3, 54) = 2.45, p = 0.073), but no interaction between the two (F (3, 54) = 0.78, p = 0.510). *Post hoc* t-tests with H-S corrections showed no statistical differences in locomotion between males and females on individual conditioning days (ps > 0.05). For 10 mg/kg, a 2×2 RM-ANOVA showed main effects of sex (F (1, 26) = 9.50, **p = 0.005) and conditioning day (F (3, 78) = 28.83, ^$$$$^p < 0.0001), but no interaction between the two (F (3, 78) = 0.50, p = 0.684). Post hoc t-tests with H-S corrections showed a sex difference in locomotion on day 1 (t(104) = 2.53, **p = 0.025), day 2 (t(104) = 2.95, *p = 0.016), and day 4 (t(104) = 2.85, *p = 0.016), with a trend on day 3 (t(104) = 1.95, p = 0.054). For 20 mg/kg, a 2×2 RM-ANOVA showed main effects of sex (F (1, 43) = 10.75, **p = 0.002) and conditioning day (F (3, 129) = 53.52, ^$$$$^p < 0.0001), but no interaction between the two (F (3, 129) = 0.35, p = 0.787). Post hoc t-tests with H-S corrections showed a sex difference in locomotion on all days (day 1: t(172) = 3.20, **p = 0.005; day 2: t(172) = 3.37, **p = 0.004; day 3: t(172) = 2.91, **p = 0.007; day 4: t(172) = 2.96, **p = 0.007). For 50 mg/kg, a 2×2 RM-ANOVA showed a trend for an effect of sex (F (1, 8) = 4.79, p = 0.060) and effect of conditioning day (F (3, 24) = 62.75, ^$$$$^p < 0.0001), as well as a trend for an interaction between the two (F (3, 24) = 2.35, p = 0.098). Post hoc t-tests with H-S corrections showed a sex difference in locomotion on day 1 (t(32) = 2.94, *p = 0.024), trend for day 2 (t(32) = 2.27, p = 0.087), and no differences on days 3 and 4 (ps > 0.70).

**Figure 3:**
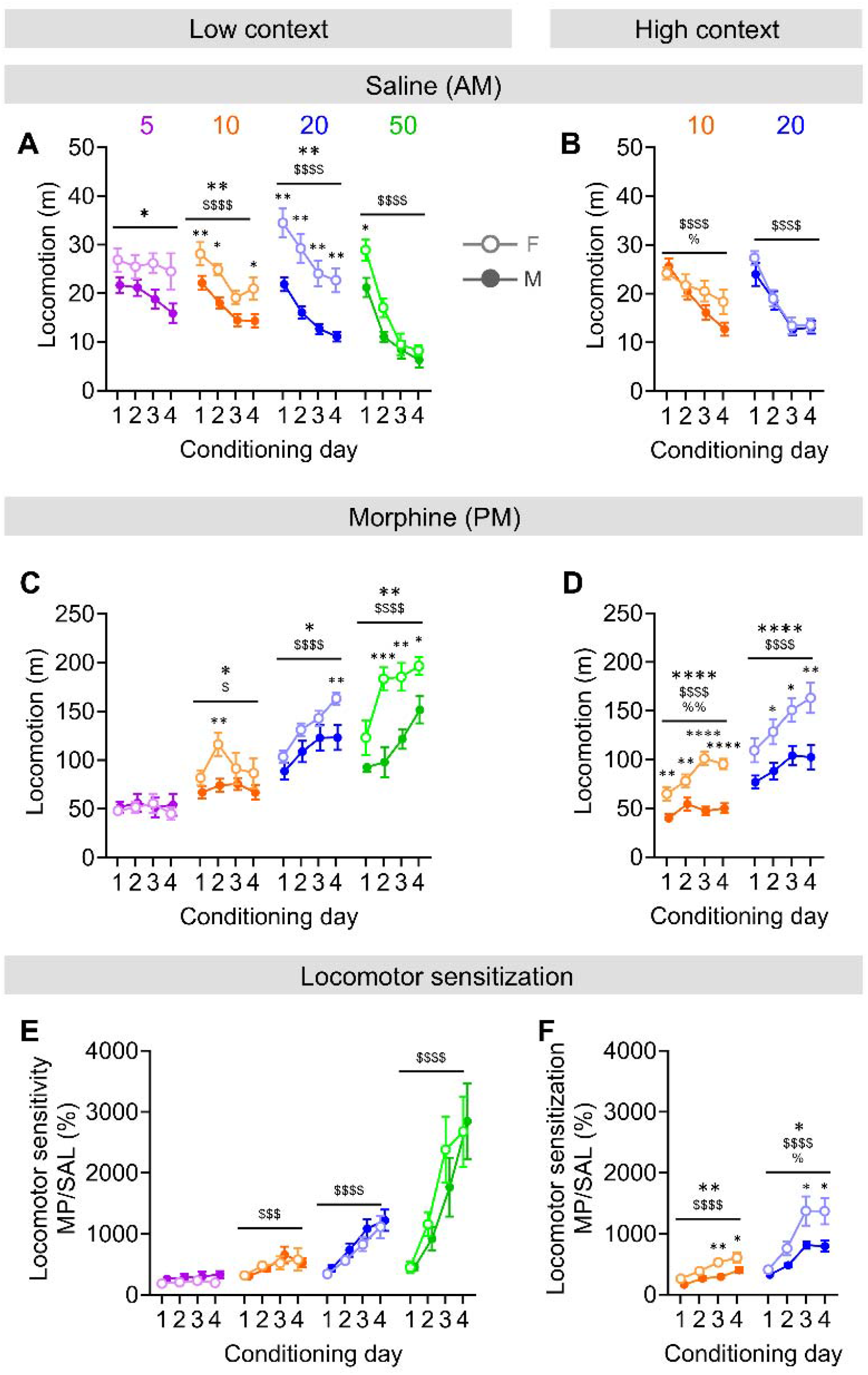
Females display greater morphine-induced locomotor sensitization than males. **A-B)** Locomotion during daily morning CPP conditioning sessions in which mice received saline vehicle injections (i.p.) before being placed in and sequestered to one compartment of the CPP apparatus in the low (**A**) or high (**B**) context versions of the assay decreased across days in both sexes. **C-D)** Locomotion during daily afternoon conditioning sessions in which mice received morphine injections (i.p.) before being place in the other compartment of the CPP apparatus in the low (**C**) and high (**D**) context versions of the assay. Locomotion increased at higher doses and to a greater extent in females compared to males. **E-F)** Locomotor sensitization to morphine across conditioning days, presented as daily locomotor sensitivity to acute morphine normalized to saline locomotion, in the low (**E**) and high (**F**) context versions of the CPP assay. Both sexes displayed dose-dependent increases in sensitization, and this was higher in females than males in the high context assay only. In 2×2 RM-ANOVAs between sex and conditioning day within dose: ^$^p < 0.05, ^$$$$^p < 0.0001 indicate main effects of day; *p < 0.05, **p < 0.01, ***p < 0.001, ****p < 0.0001 indicate main effect of sex, as well as in *post hoc* t-tests between males and females within day; ^%^p < 0.05 indicates sex x day interaction. Effects from 2×2 RM-ANOVAs between dose and day within sex are not indicated in figure but are reported in the text.

We further evaluated whether saline locomotion across days depended on the morphine dose administered during conditioning (statistics below are not indicated in Fig. 3A). For females, a 2×2 RM-ANOVA showed no effect of morphine dose (F (3, 52) = 1.74, p = 0.171) but an effect of conditioning day (F (3, 156) = 37.79, p < 0.0001) and an interaction between the two (F (9, 156) = 4.42, p < 0.0001). Post hoc comparisons with H-S corrections showed that females did not show changes in locomotion on saline sessions in the 5 mg/kg morphine mice (all ps > 0.85) but did have decreased locomotion across days in the 10 mg/kg (day 1 vs. days 3 and 4: ps < 005; day 2 vs. day 3: p < 0.05; all other comparisons: ps > 0.15), 20 mg/kg (day 3 vs. 4: p > 0.25; all other comparisons: ps < 0.0005), and 50 mg/kg (day 3 vs. 4: p > 0.687; all other comparisons: ps < 0.05) dose groups). In contrast, a 2×2 RM-ANOVA in males showed main effects of morphine concentration (F (3, 43) = 4.33, p = 0.009) and conditioning day (F (3, 129) = 59.70, p < 0.0001), as well as an interaction between the two (F (9, 129) = 2.79, p = 0.005). Post hoc comparisons with H-S corrections showed that locomotion differed between the 5 mg/kg and 50 mg/kg doses (p < 0.010), but no other comparisons were significantly different (ps > 0.05). In addition, males displayed reduced locomotion across saline sessions within each morphine dose group (5 mg/kg: days 1 and 2 vs. day 4: ps < 0.005; all others: ps > 0.20; 10 mg/kg: day 3 vs 4: p > 0.90; all other ps < 0.01; 20 mg/kg: day 3 vs 4: p > 0.15; all other ps < 0.05; 50 mg/kg: day 1 vs. all other days: ps < 0.0001; all other comparisons: ps > 0.05). Overall, these results show that while females have higher saline locomotion (similar to basal locomotion on pretest), both sexes displayed a decrease in saline locomotion across conditioning days that became more robust across increasing morphine doses. These results suggest that mice may be in withdrawal during daily saline sessions, particularly in groups given high morphine doses.

Interestingly, the effects of sex and morphine dose on saline locomotion were less robust in the high context version of the CPP assay (Fig. 3B). In a 2×2 RM-ANOVA between sex and day for 10 mg/kg, there was no effect of sex (F (1, 19) = 1.37, p = 0.256) but an effect of conditioning day (F (3, 55) = 21.03, ^$$$$^p < 0.0001) and an interaction between the two (F (3, 55) = 2.932, ^%^p = 0.041). For 20 mg/kg, a 2×2 RM-ANOVA showed no effect of sex (F (1, 20) = 0.33, p = 0.572) but an effect of conditioning day (F (3, 60) = 85.61, ^$$$$^p < 0.0001) and no interaction between the two (F (3, 60) = 1.13, p = 0.344). We further analyzed locomotion across conditioning days and increasing doses of morphine within each sex showed that saline locomotion (statistics below are not indicated in Fig. 3B). A 2×2 RM mixed model in females showed no effect of morphine dose (F (1, 21) = 2.04, p = 0.168) but an effect of conditioning day (F (3, 61) = 24.27, p < 0.0001) and an interaction between the two (F (3, 61) = 5.27, p = 0.003). Post hoc comparisons with H-S corrections showed that females had decreased locomotion across days that was more robust for the 20 mg/kg group than 10 mg/kg group (10 mg/kg: day 1 vs. day 4: p <0.05, all other comparisons: ps > 0.15; 20 mg/kg: day 3 vs. day 4: p > 0.90, all other ps < 0.01), but locomotion between doses was only different on day 3 (p = 0.028; all other ps > 0.15). For males, a 2×2 RM-ANOVA showed no effect of morphine dose (F (1, 18) = 0.58, p = 0.456) but an effect of conditioning day (F (3, 54) = 98.15, p < 0.0001) and no interaction between the two (F (3, 54) = 2.02, p = 0.122). Post hoc comparisons with H-S corrections showed that locomotion decreased across days for both doses (20 mg/kg day 3 vs. 4: p > 0.70; all other comparisons ps < 0.005).

Females also had higher morphine-induced hyperlocomotion than males across increasing doses in the low context assay (Fig. 3C). A 2×2 RM-ANOVA for the 5 mg/kg dose of morphine showed no effects of sex, conditioning day, or interaction between the two (ps > 0.55). For 10 mg/kg, a 2×2 RM-ANOVA showed main effects of sex (F (1, 26) = 4.52, *p = 0.043) and conditioning day (F (3, 78) = 3.97, ^$^p = 0.010), but no interaction between the two (F (3, 78) = 1.885, p = 0.139). Post hoc t-tests with H-S corrections showed a sex difference in locomotion on day 2 (t(104) = 3.10, **p = 0.010) but no other days (ps > 0.35). For 20 mg/kg, a 2×2 RM-ANOVA showed main effects of sex (F (1, 43) = 5.494, *p = 0.024) and conditioning day (F (3, 129) = 29.01, ^$$$$^p < 0.0001), but no interaction between the two (F (3, 129) = 2.089, p = 0.105). Post hoc t-tests with H-S corrections showed a sex difference in locomotion on day 4 (t(172) = 3.26, **p = 0.005) but no other days (ps > 0.15). And, for 50 mg/kg, a 2×2 RM-ANOVA showed main effects of sex (F (1, 8) = 24.31, **p = 0.001) and conditioning day (F (3, 24) = 12.03, ^$$$$^p < 0.0001), but no interaction between the two (F (3, 24) = 2.224, p = 0.111). Post hoc t-tests with H-S corrections showed a trend for a sex difference in locomotion on day 1 (t(32) = 1.70, p = 0.098) and significant difference on days 2 (t(32) = 4.77, ***p = 0.0002), 3 (t(32) = 3.55, **p = 0.004), and 4 (t(32) = 2.50, *p = 0.035). We further evaluated the locomotor effects of morphine during conditioning within sex across increasing doses (statistics below are not indicated in Fig. 3C). For females, a 2×2 RM-ANOVA showed main effects of morphine concentration (F (3, 52) = 31.71, p < 0.0001) and conditioning day (F (3, 156) = 15.64, p < 0.0001), as well as an interaction between the two (F (9, 156) = 6.10, p = 0.0002). Post hoc comparisons with H-S corrections confirmed that locomotion increased across all doses of morphine, as each increasing dose elicited greater locomotion than all other lower doses (all ps ≤ 0.001). In addition, females did not display a sensitized response to morphine at lower doses (5 mg/kg: all ps > 0.90; 10 mg/kg: day 2 vs. day 1 (t(156) = 3.38, p = 0.005; day 2 vs. day 4 (t(156) = 2.90, p = 0.021; all other ps > 0.05), but robust sensitization emerged at 20 mg/kg (except day 2 vs. 3, all ps < 0.005) and was reached by day 2 for 50 mg/kg (day 1 vs all others: ps < 0.001; all other ps > 0.75). Similarly in males, a 2×2 RM-ANOVA showed main effects of morphine concentration (F (3, 43) = 10.54, p < 0.0001) and conditioning day (F (3, 129) = 9.933, p < 0.0001), as well as an interaction between the two (F (9, 129) = 3.920, p < 0.0001). Post hoc comparisons with H-S corrections confirmed that locomotion increased across morphine concentrations, plateauing at 20 mg/kg, as 5 vs. 10 mg/kg and 20 vs. 50 mg/kg did not significantly differ from one another (ps > 0.25) but all other dose comparisons did (ps ≤ 0.002). As in females, males did not display locomotor sensitization across days of conditioning at lower doses (5 and 10 mg/kg: all ps > 0.75) but robust sensitization emerged at 20 mg/kg (day 1 vs all other days: ps < 0.05; all other comparisons ps > 0.10) and 50 mg/kg (days 1 and 2 vs day 4: ps < 0.0002; all other ps > 0.05).

Morphine-induced hyperlocomotion was even more robust in the high context CPP assay than in the low context version of the assay, with females showing even greater locomotor responses to acute and repeated daily morphine exposure than males (Fig. 3D). A 2×2 RM-ANOVA in the high context showed main effects of sex (F (1, 19) = 34.97, ****p < 0.0001) and conditioning day (F (3, 57) = 9.16, ^$$$$^p < 0.0001) and an interaction between the two (F (3, 57) = 5.22, ^%%^p = 0.003). Post hoc t-tests with H-S corrections showed a sex difference in locomotion on all days (day 1: (t(76) = 2.95, **p = 0.009; day 2: t(76) = 2.75, **p = 0.009; day 3: t(76) = 6.36, ****p < 0.0001; day 4: t(76) = 5.36, ****p < 0.0001). Similarly, for 20 mg/kg, a 2×2 RM-ANOVA showed main effects of sex (F (1, 20) = 7.942, *p = 0.011) and conditioning day (F (3, 60) = 21.32, ^$$$$^p < 0.0001) but not an interaction between the two (F (3, 60) = 2.274, p = 0.089). Post hoc t-tests with H-S corrections showed a trend for a sex difference in locomotion on day 1 (t(80) = 1.85, p = 0.067) and significant differences on all other days (day 2: t(80) = 2.31, *p = 0.047; day 3: t(80) = 2.67, *p = 0.027; day 4: t(80) = 3.50, **p = 0.003). Evaluation of the locomotor effects of morphine within sex across doses in the high context confirmed that females displayed greater sensitization to 10 mg/kg morphine in the high context assay than on the low context version (statistics below are not indicated in Fig. 3D). A 2×2 RM-ANOVA in females showed main effects of morphine concentration (F (1, 21) = 14.75, p = 0.001) and conditioning day (F (3, 63) = 29.21, p < 0.0001), but no interaction (F (3, 63) = 1.90, p = 0.139). Post hoc comparisons with H-S corrections confirmed that locomotion increased across days for 10 mg/kg (day 1 vs. days 3 and 4: ps < 0.001; day 2 vs. day 3: p = 0.011; all others: ps > 0.05), and 20 mg/kg (day 1 vs. all others and day 2 vs. all others: ps < 0.05; days 3 vs. 4: p > 0.05). In addition, locomotion was higher for 20 mg/kg than 10 mg/kg for all days (ps < 0.005). For males, locomotor sensitization was more robust for 20 mg/kg than 10 mg/kg, just as we observed in the low context assay. A 2×2 RM-ANOVA showed main effects of morphine concentration (F (1, 18) = 24.57, p = 0.0001) and conditioning day (F (3, 54) = 5.49, p = 0.002), and a trend for interaction (F (3, 54) = 2.53, p = 0.067). Post hoc comparisons with H-S corrections showed that locomotion increased across days for 20 mg/kg (day 1 vs. days 3 and 4: ps < 0.005; all others: ps > 0.10) but not 10 mg/kg (all ps > 0.20). In addition, locomotion was higher for 20 mg/kg than 10 mg/kg for all days (ps < 0.005).

We examined the locomotor sensitivity to acute morphine, calculated as the distance traveled during the PM morphine session normalized to distance traveled during the AM morphine session each day (e.g., C1 sensitivity (%) = C1 morphine distance / C1 saline distance), in order to evaluate the change in this relationship across conditioning days, a term called locomotor sensitization (**Fig. E,F**). In the low context assay, we found that due to the proportionally larger increases in females than males in locomotion during morphine sessions (Fig. 3C) and decreases in locomotion during saline sessions (Fig. 3A), resulting locomotor sensitization was similar between sexes (Fig. 3E). A 2×2 RM-ANOVA for the 5 mg/kg dose showed no effect of sex or conditioning day or an interaction (ps > 0.15). For 10 mg/kg, a 2×2 RM-ANOVA showed an effect of conditioning day (F (3, 78) = 6.47, ^$$$^p = 0.0006) but no effect of sex (F (1, 26) = 0.00, p = 0.997) or interaction (F (3, 78) = 0.79, p = 0.502). Similarly for 20 mg/kg, there was an effect of conditioning day (F (3, 129) = 30.70, ^$$$$^p < 0.0001) but no effect of sex (F (1, 43) = 1.15, p = 0.289) or interaction (F (3, 129) = 0.37, p = 0.775), and for 50 mg/ kg, there was an effect of conditioning day (F (3, 24) = 23.29, ^$$$$^p < 0.0001) but no effect of sex (F (1, 8) = 0.15, p = 0.707) or interaction (F (3, 24) = 0.63, p = 0.602). Examination of locomotor sensitization within sex showed a dose-dependent increase in locomotor sensitization in both females and males (statistics below are not indicated in Fig. 3E). For females, a 2×2 RM-ANOVA showed effects of morphine dose (F (3, 52) = 14.57, p < 0.0001) and conditioning day (F (3, 156) = 34.20, p < 0.0001), as well as an interaction between the two (F (9, 156) = 8.98, p < 0.0001). Post hoc comparisons with H-S corrections showed that females did not show sensitization at 5 or 10 mg/kg doses of morphine (all day comparisons: ps > 0.45), but they did show locomotor sensitization at 20 mg/kg (all ps < 0.05) and 50 mg/kg (day 3 vs day 4: p > 0.20; all other ps ≤ 0.010). In addition, sensitization increased across morphine doses, as 5 vs. 10 mg/kg and 10 vs. 20 mg/kg did not significantly differ (ps > 0.15) but all other dose comparisons did (ps < 0.010). For males, a 2×2 RM-ANOVA showed effects of morphine dose (F (3, 43) = 14.38, p < 0.0001) and conditioning day (F (3, 129) = 56.31, p < 0.0001), as well as an interaction between the two (F (9, 129) = 14.43, p < 0.0001). Post hoc comparisons with H-S corrections showed that males did not show sensitization at 5 mg/kg (all day comparisons ps > 0.95) or 10 mg/kg (day 1 vs day 3: p < 0.05; but all other day comparisons: ps > 0.20). However, they did show locomotor sensitization at 20 mg/kg (day 3 vs. day 4: p > 0.20; all other ps < 0.05) and 50 mg/kg (all day comparisons ps < 0.05).

In the high context version of the assay, the more robust locomotor response to morphine in females than males (Fig. 3D) but lack of substantial sex difference in saline locomotion (Fig. 3B) resulted in greater locomotor sensitization in females than males (Fig. 3F). For 10 mg/kg, a 2×2 RM mixed model showed main effects of sex (F (1, 19) = 10.66, **p = 0.004) and conditioning day (F (3, 55) = 24.25, ^$$$$^p < 0.0001) but no interaction between the two (F (3, 55) = 1.772, p = 0.163). Post hoc t-tests with H-S corrections showed a sex difference in locomotor sensitization on days 3 and 4 but not days 1 or 2 (day 1: (t(74) = 1.18, p = 0.240; day 2: t(74) = 1.78, p = 0.153; day 3: t(74) = 3.55, **p = 0.003; day 4: t(74) = 3.03, *p = 0.010). Similarly, at 20 mg/kg, a 2×2 RM-ANOVA showed main effects of sex (F (1, 20) = 4.92, *p = 0.038) and conditioning day (F (3, 60) = 29.72, ^$$$$^p < 0.0001) and an interaction between the two (F (3, 60) = 3.08, ^%^p = 0.034). Post hoc t-tests with H-S corrections showed a sex difference in locomotor sensitization on days 3 and 4 but not days 1 or 2 (day 1: (t(80) = 0.36, p = 0.720; day 2: t(80) = 1.35, p = 0.329; day 3: t(80) = 2.73, *p = 0.026; day 4: t(80) = 2.80, *p = 0.026).

We further evaluated locomotor sensitization within sex across the two doses in the high context assay (statistics below are not indicated in Fig. 3F). For females, a 2×2 RM mixed model showed main effects of morphine concentration (F (1, 21) = 12.14, p = 0.002) and conditioning day (F (3, 61) = 23.15, p < 0.0001) and an interaction (F (3, 61) = 23.15, p < 0.0001). Post hoc comparisons with H-S corrections confirmed that locomotion increased across days for 20 mg/kg (day 3 vs 4: p > 0.95, all other ps < 0.05) but not 10 mg/kg (all ps > 0.05). In addition, locomotion was higher for 20 mg/kg than 10 mg/kg for days 3 and 4 (ps < 0.0005) but not days 1 or 2 (ps > 0.10). For males, a 2×2 RM-ANOVA showed main effects of morphine concentration (F (1, 18) = 42.57, p < 0.0001) and conditioning day (F (3, 54) = 39.32, p < 0.0001), as well as an interaction between the two (F (3, 54) = 9.82, p < 0.0001). Post hoc comparisons with H-S corrections showed that locomotion increased slightly across the four days of conditioning in the 10 mg/kg dose group (day 1 vs. day 4: p < 0.001; all other ps > 0.05) and robustly in the 20 mg/kg group (day 3 vs. day 4: p > 0.75, all other ps < 0.05). In addition, locomotion was higher for 20 mg/kg than 10 mg/kg for all days (ps < 0.05).

### 3.4 Opposing relationships between the locomotor effects of morphine and CPP in females and males

We next assessed the magnitude of conditioned place preference (CPP) for the morphine-paired compartment of the CPP apparatus on post-conditioning day 1 (P1) in the low and high context versions of the assay across morphine doses and evaluated whether it was associated with the locomotor effects of morphine. We found that neither male nor female mice expressed a CPP one day following conditioning in the low context assay for any dose of morphine (Fig. 4A), even though 10, 20, and 50 mg/kg doses were sufficient to induce robust locomotor effects suggesting ample bioavailability of the drug (Figs. 2 and 3). A 2×2 ANOVA in the low context assay show no effect of sex or morphine concentration (ps > 0.40), and one-sample t-tests showed that neither males nor females showed a significant CPP at any dose (ps > 0.20). In contrast, both female and male mice expressed a CPP in the high context version (Fig. 4B). A 2×2 ANOVA also showed no effect of sex of morphine concentration (ps 0.35), however one-sample t-tests showed that females acquired a CPP at both the 10 and 20 mg/kg doses (10 mg/kg: t(10) = 3.92, ^##^p = 0.003; 20 mg/kg: t(12) = 2.58, ^#^p = 0.024) and males display a P1 CPP for 10 mg/kg (t(9) = 3.66, ^##^p = 0.005) but not 20 mg/kg (t(8) = 1.68, p = 0.132). Analysis of the relationship between the locomotor effects of morphine and CPP score in the high context assay revealed sex-dependent and opposing relationships (Fig. 4C**,D**).

**Figure 4:**
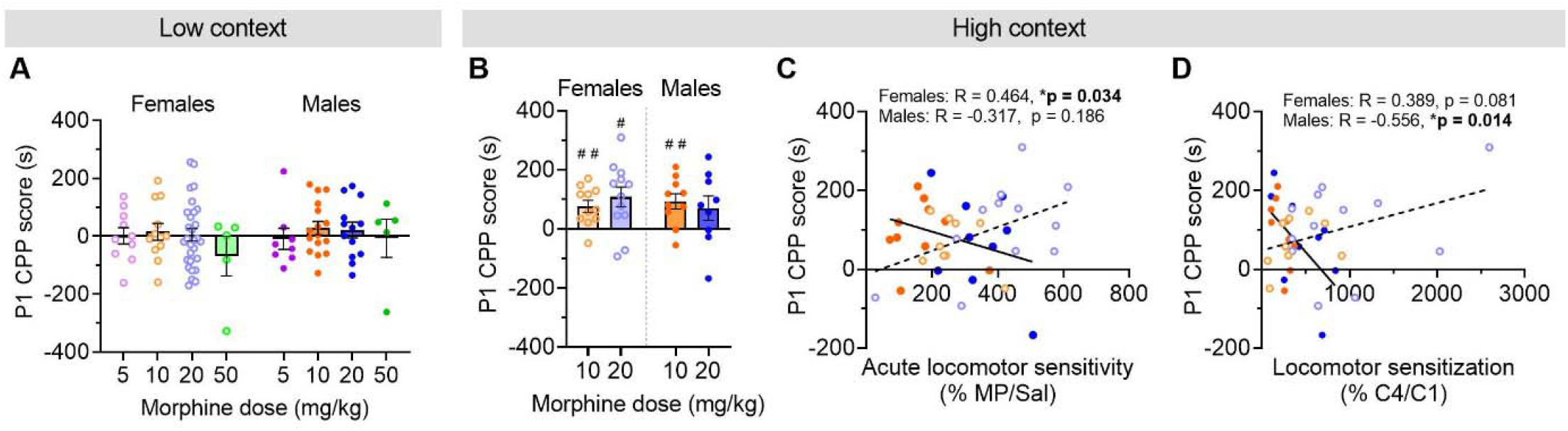
Sex-dependent relationship between the locomotor effects of morphine and morphine-context association strength. **A-B)** Conditioned place preference (CPP) scores on post-conditioning day 1 (P1) on the low (**A**) and high (**B**) context CPP assays showing that males and females displayed a CPP following conditioning in the high context, but not low context, assay. ^#^p < 0.05, ^##^p < 0.01 indicate group CPP scores significantly higher than the null hypothesis of 0 s difference between P1 and pretest preference for the morphine-paired compartment. **C)** There was positive correlation between acute locomotor sensitivity to morphine on C1 and CPP on P1 in females but no association in males. **D)** There was a trend for a positive correlation between locomotor sensitization to morphine (C4 sensitivity / C1 sensitivity) and CPP in females, but a significant negative correlation in males. *p < 0.05 for Pearson’s correlations.

P1 CPP score was significantly positively correlated with acute locomotor sensitivity (from Fig. 2) in females (Y = 0.301x – 13.31; R = 0.464, *p = 0.034), while there was no relationship in males (Y = −0.250x + 145.1; R = −0.317, p = 0.186; Fig. 4C). In contrast, male P1 CPP was significantly negatively correlated with locomotor sensitization (from Fig. 3; Y = −0.253x + 170.2; R = −0.556, *p = 0.014), while female CPP showed a trend for positive correlation with locomotor sensitization (Y = 0.060x + 48.03; R = 0.389, p = 0.081; Fig. 4D). These surprising results suggest that the locomotor effects of morphine bidirectionally predict CPP in males and females.

### 3.5 Sex-dependent relationships between morphine withdrawal and CPP expression in females and males

We found that during CPP conditioning, locomotion during the morning saline sessions decreased more dramatically across days in a dose-dependent manner in both sexes and contexts (Fig. 3A**,B**), suggesting that mice may have been in morphine withdrawal during those sessions. Therefore, we examined locomotion across the pre- and post-tests on P1, as well as on an additional test 10 days later into home cage abstinence on P11, within sex to further investigate the dose-dependent hypolocomotive phenotype we observed during conditioning. We first examined this in the low context assay to evaluate these locomotor effects independent of morphine-context learning, as mice did not establish a CPP in this version of the assay (Fig. 4). In females (Fig. 5A), a 2×2 RM-ANOVA showed a main effect of test day (F (2, 102) = 45.87, ****p < 0.0001) and no effect of morphine dose (F (3, 51) = 1.448, p = 0.240) or interaction (F (6, 102) = 0.9939, p = 0.434). Post hoc comparisons with H-S corrections between tests within dose showed that P1 locomotion was lower than pretest for all doses (all ^$$$^ps ≤ 0.0005) and lower than P11 for all doses but 5 mg/kg (5 mg/kg: p > 0.05; all other ^&^ps < 0.05).

**Figure 5:**
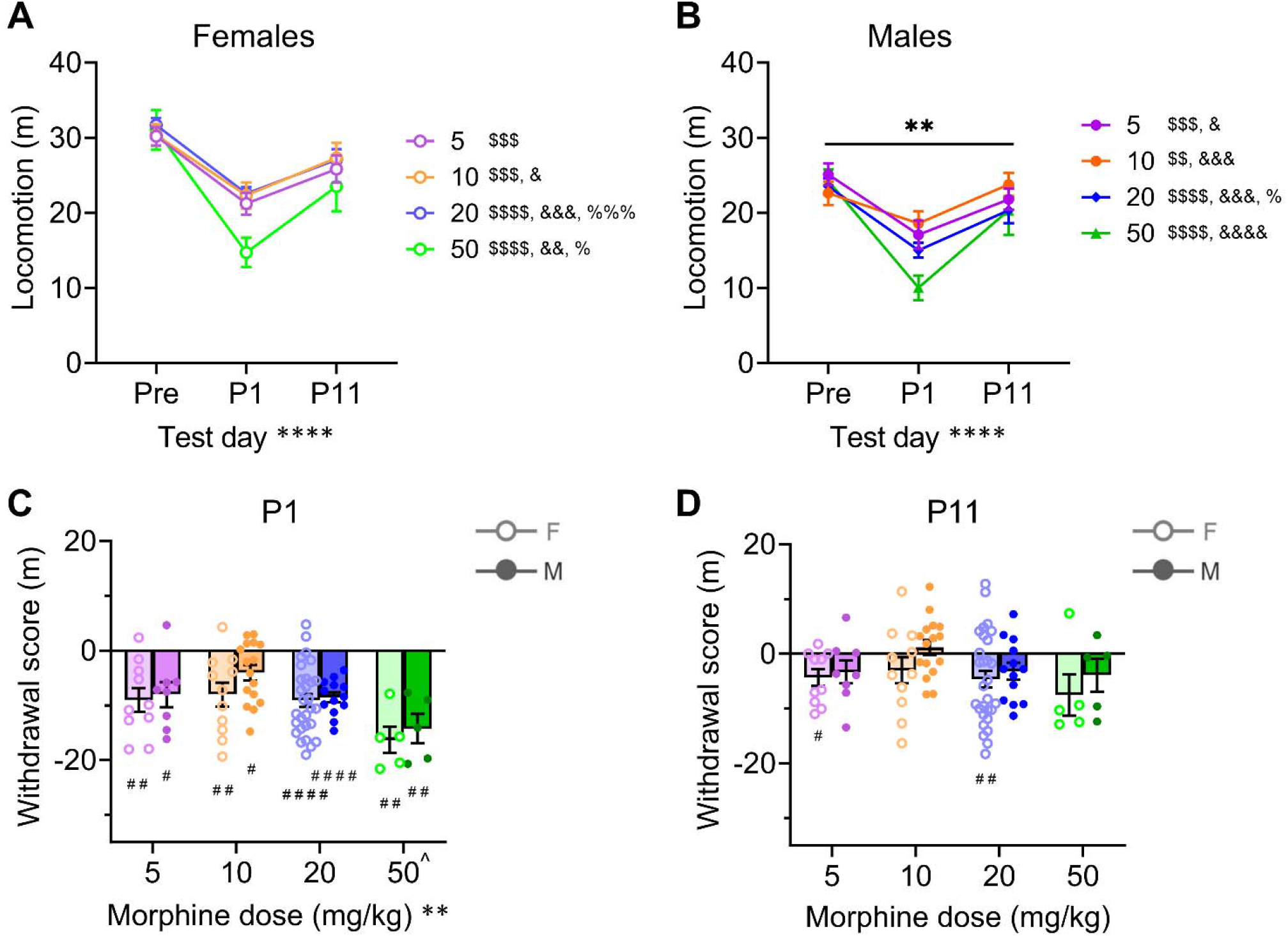
Mice display a dose-dependent withdrawal-induced hypolocomotion on post-conditioning day 1 that recovers 10 days later in the low context version of the CPP assay. **A-B)** Locomotion on pretest, P1 test, and P11 test in females (**A**) and males (**B**) administered different doses of morphine during CPP conditioning, showing a hypolocomotive phenotype on P1 across doses in both sexes. **p < 0.01, ****p < 0.0001 indicate main effect of test day and interaction between dose and test day in 2×2 RM-ANOVAs; ^$$^p < 0.01, ^$$$^p < 0.001, ^$$$$^p < 0.0001 indicate differences between pretest and P1, ^&^p < 0.05, ^&&^p < 0.01, ^&&&^p < 0.001, ^&&&&^p < 0.0001 indicate differences between P1 and P11, and ^%^p < 0.05. ^%%%^p < 0.001 indicate differences between pretest and P11. **C-D)** Withdrawal scores, calculated as test locomotion minus pretest locomotion, on P1 (**C**) and P11 (**D**) in males and females. **p < 0.01 indicates a main effect of morphine dose in 2×2 RM-ANOVA between dose and sex; ^p < 0.05 indicate *post hoc* t-test differences between 50 mg/kg dose and all other doses; ^#^p < 0.05, ^##^p < 0.01, ^####^p < 0.0001 in one-sample t-tests compared to the null hypothesis of 0 within group.

P11 locomotion was lower than pretest for 20 and 50 mg/kg (^%^ps < 0.05) but not 5 or 10 mg/kg (ps > 0.05). In males (Fig. 5B), a 2×2 RM-ANOVA showed no effect of morphine concentration (F (3, 38) = 1.058, p = 0.378), but did show a main effect of test day (F (2, 76) = 52.72, ****p < 0.0001) and interaction between dose and test (F (6, 76) = 3.184, **p = 0.008). Post hoc comparisons with H-S corrections between tests within dose showed that P1 locomotion was lower than pretest for all doses (all ^$$^ps ≤ 0.005) and lower than P11 for all doses (^&^ps < 0.05). P11 locomotion was lower than pretest for 20 mg/kg (^%^p < 0.05) but not the other doses (ps > 0.05). Together these data show that P1 locomotion was reduced at P1 and at least partially recovered by P11.

We explored whether there were sex differences in the magnitude of locomotion change for each post-test across doses using withdrawal locomotion scores (test locomotion – pretest locomotion). We found that for P1 (Fig. 5C), a 2×2 RM-ANOVA showed a main effect of morphine dose (F (3, 89) = 5.734, **p = 0.001) but no effect of sex (F (1, 89) = 1.816, p = 0.181) or interaction (F (3, 89) = 0.4414, p = 0.724). Post hoc comparisons with H-S corrections across morphine doses regardless of sex showed that the P1 withdrawal locomotion scores for 50 mg/kg were significantly greater than for all other doses (^ps < 0.05), while no other doses differed from each other (ps > 0.20). In a similar analysis for P11 (Fig. 5D), there were no effects of morphine dose or sex or an interaction between the two variables (ps > 0.10). One-sample t-tests for each dose x sex x test day group showed that there was a hypolocomotive phenotype on P1 for all groups (^#^ps < 0.05) but only for 5 and 20 mg/kg in females on P11 (^#^ps < 0.05; all other ps > 0.05). Altogether, locomotion data from the test days in the low context assay, in which CPP was not displayed, revealed that there was a dose dependent hypolocomotion on P1 that recovered, at least partially, by P11. These results are consistent with our conditioning data showing that hypolocomotion during daily saline conditioning sessions became exacerbated across increasing doses of morphine (Fig. 3), a sign of withdrawal.

In the high context version of the CPP assay, we tested mice approximately every 10 days until post-conditioning day 70 (P70) and again at P100 to examine the stability of the morphine-context association, as well as its relationship to locomotor behavior on the test. We found that CPP score in females, which was significant on P1 (Fig. 4B), was stable across the post-tests; a 2×2 RM-ANOVA showed no effect of test day (F (8, 168) = 1.47, p = 0.172) or morphine dose (F (1, 21) = 0.73, p = 0.403) or interaction between the two (F (8, 168) = 0.35, p = 0.943) on CPP score across all tests (Fig. 6A). In contrast, this analysis in males (Fig. 6B) showed a main effect of test day (F (8, 136) = 3.34, **p = 0.002) with no effect of morphine dose (F (1, 17) = 0.29, p = 0.598) or interaction (F (8, 136) = 0.49, p = 0.865). Post hoc analysis of CPP score across the first five post-tests showed that male CPP increased from P1 to P11, followed by a slow decay in CPP across P21 to P38. Direct comparisons of CPP between test days with H-S corrections for multiple comparisons showed that P11 and P21 CPP were significantly higher than P1 (^$^ps < 0.05) and P38 (^##^ps < 0.01), while no other comparisons were different (ps > 0.20). Given the change in CPP across P1 and P11 in males but not females, we examined whether these two CPP tests had a different relationship in males and females (Fig. 6C). Pearson’s correlation analyses showed that P1 and P11 CPP were highly positively correlated with one another in females (Y = 0.704x + 45.67; R = 0.828, ****p < 0.0001) but not associated in males (Y = 0.257x + 144.0; R = 0.314, p = 0.190).

**Figure 6:**
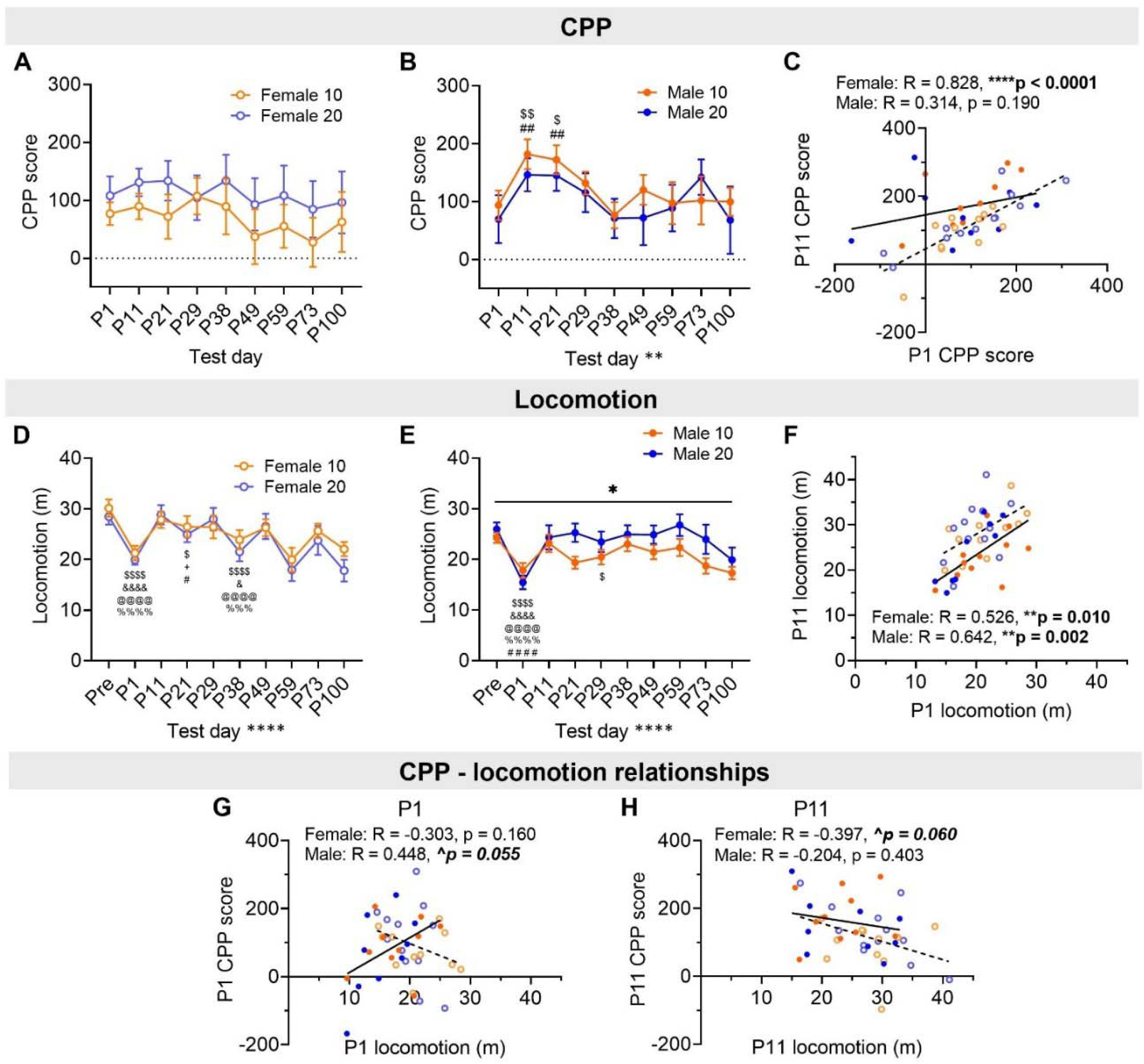
Sex-dependent CPP and CPP-locomotion relationships across abstinence in the high context assay. **A-B)** Conditioned place preference (CPP) scores in females (**A**) and males (**B**) across repeated testing, showing sex-dependent changes over time. **C)** Correlations between P1 and P11 CPP scores, showing a significant positive association between P1 and P11 CPP in females but not males. **D-E)** Locomotion during CPP tests in females (**D**) and males (**E**), showing a hypolocomotive phenotype on P1. **F)** Correlations between P1 and P11 CPP scores in females and males, demonstrating a positive correlation between P1 and P11 locomotion in both sexes. **G-H)** Correlations between locomotion and CPP scores on P1 (**G**) and P11 (**H**), illustrating opposing relationships in females and males. *p < 0.05, **p < 0.01, ****p < 0.0001 indicate a main effect of test day or interaction between test day and morphine dose in 2×2 RM-ANOVA or Pearson’s correlation; ^$^p < 0.05, ^$$^p < 0.01, ^$$$$^p < 0.0001 indicate difference from pretest, ^+^p < 0.05 from P1, ^&^p < 0.05, ^&&&&^p < 0.0001 from P11, ^@@@@^p < 0.0001 from P21, ^%%%%^p < 0.0001 from P29, ^#^p < 0.05 from P38 in *post hoc* t-tests following 2×2 RM-ANOVAs comparing the first five tests.

We also evaluated locomotion across high context CPP tests, finding that as in the low context version of the assay, mice displayed a hypolocomotive phenotype on P1 that recovered by P11 (Fig. 6D**,E**). In females (Fig. 6D), a 2×2 RM-ANOVA showed a main effect of test day (F (9, 189) = 12.66, ****p < 0.0001) with no effect of morphine dose (F (1, 21) = 0.40, p = 0.535) or interaction (F (9, 189) = 0.69, p = 0.713). Direct comparisons between locomotion on the first five CPP tests with H-S corrections for multiple comparisons showed that locomotion was lower on P1 than on the pretest but then rebounded on P11 before decaying between P21 and P38. Specifically, locomotion on P1 and P38 were lower than on all other tests (ps < 0.05 as indicated) but not different from each other (p > 0.20), and P21 locomotion was different from pretest, P1, and P38 (ps < 0.05 as indicated) but not P11 or P29 (ps > 0.05).

In males (Fig. 6E), a 2×2 RM-ANOVA showed a main effect of test day (F (9, 153) = 11.07, ****p < 0.0001) with no effect of morphine dose (F (1, 17) = 2.010, p = 0.174) but an interaction between the two (F (9, 153) = 2.12, *p = 0.031). Direct comparisons with H-S corrections showed no differences between 10 mg/kg and 20 mg/kg doses within individual test days (ps > 0.15), but direct comparisons between locomotion on the first five CPP tests showed P1 locomotion lower than all other test days (ps < 0.0001 as indicated) and P29 lower than pretest locomotion (^$^p < 0.05). Taken together, we found that locomotion was suppressed on P1 and rebounded on P11 in both sexes; this was followed by a slow decay in locomotion over subsequent tests in females but not males. While both sexes showed a hypolocomotive phenotype on P1 that rebounded by P11, there was a positive correlation between locomotion on the two tests in both sexes (females: Y = 0.778x + 12.36; R = 0.526, **p = 0.01; males: Y = 0.888x + 8.91; R = 0.642, **p = 0.002; Fig. 6F). Given the sex-dependent nature of locomotion and CPP changes over time, we evaluated whether there were sex-dependent relationships between locomotion and CPP. On P1 (Fig. 6G), there was a trend for a positive correlation between locomotion and CPP (Y = 10.24*X − 88.72; R = 0.448, p = 0.055) but a nonsignificant negative association in females (Y = −7.08x + 238.8; R = −0.303, p = 0.160). On P11, this negative association in females became stronger into a trend (Y = −5.33x + 262.6; R = −0.397, p = 0.60), and the positive trend in males was ablated (Y = −2.749x + 230.3; P = −0.204, p = 0.403). These data suggest that withdrawal-induced hypolocomotion may interfere with the expression of CPP on P1 in males; in contrast, morphine sensitivity may promote CPP in females.

### 3.6 Effects of contextual richness on the relationships between the locomotor effects of morphine and morphine-context association strength

Altogether, our locomotor data suggest that mice are in withdrawal during saline conditioning and P1 test sessions (Fig. 3A**,B**; Fig. 6D**,E**) and that the effects of this are sex-dependent and affect the expression of CPP. Overall, acute morphine sensitivity and locomotor sensitization were predictive of higher CPP in females and lower CPP in males in the high context version of the assay, while test day withdrawal-induced hypolocomotion negatively predicted CPP in males (summarized in Fig. 7A). Interestingly, while CPP was not achieved within any groups in the low context version of the assay (Fig. 4A), we observed weaker but similar relationships between these variables as observed in the high context in females but not males (summarized in Fig. 7B). This suggests that while context information was insufficient to support CPP overall in the low context version of the assay, some learning may have occurred in a subset of female mice, and this was positively correlated with the locomotor effects of morphine.

**Figure 7:**
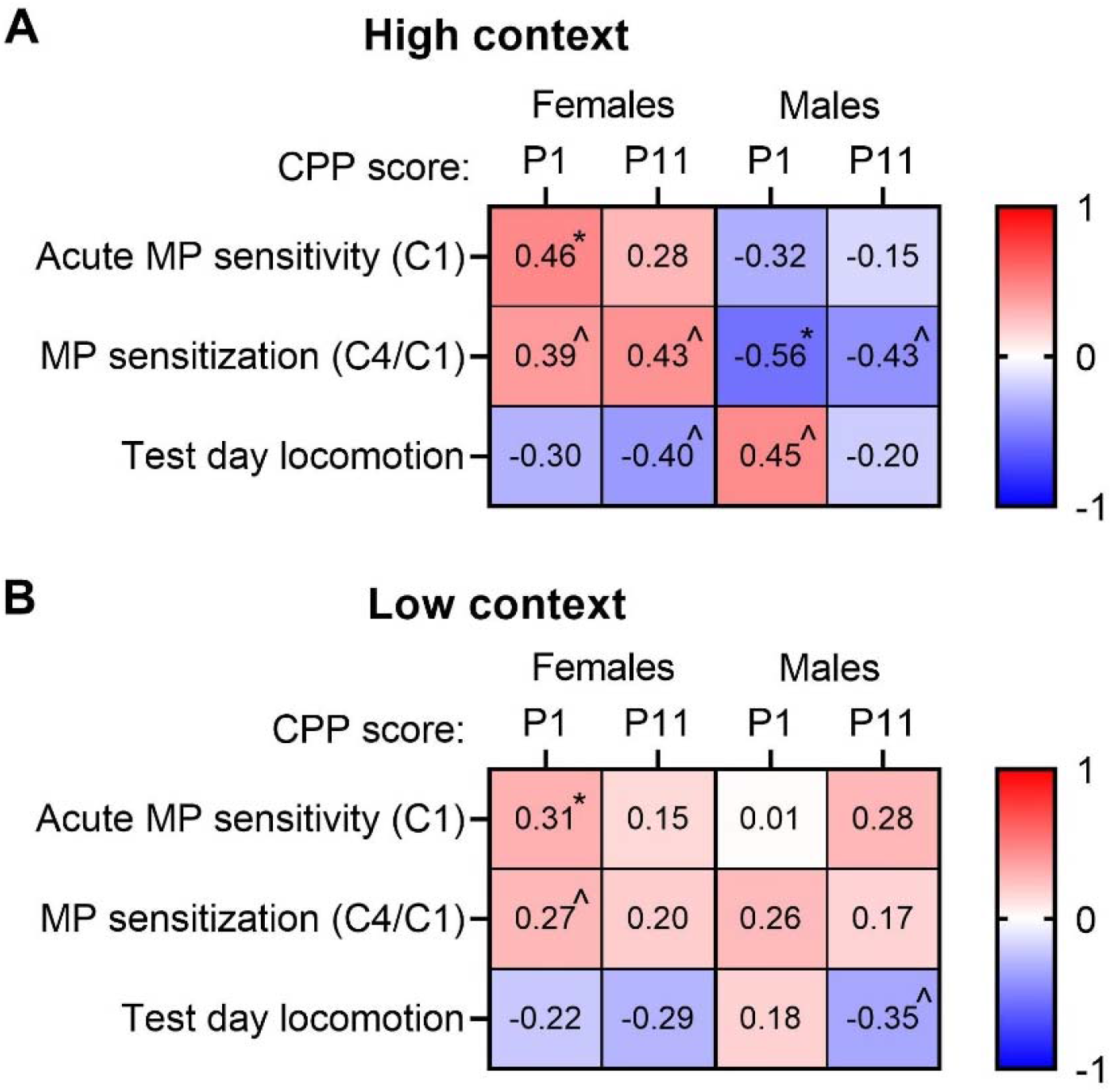
Females and males display opposing relationships between morphine- and withdrawal-induced locomotion and CPP scores. Heat map correlation matrices between CPP scores and locomotor measures in the high (**A**) and low (**B**) context versions of the CPP assay, with R values presented. *p < 0.05, ^p < 0.10 indicate significance of Pearson’s correlations.

## 4 Discussion

The development and persistence of OUD involves the formation and stability of learned associations between opioids and the context in which they are experienced, and the CPP paradigm offers a useful tool for specifically studying drug-context associations and the variables that modulate their strength and stability (Bardo & Bevins, 2000; Bardo *et al*., 2015). As females have been largely omitted from the literature, here we examined the extent to which these factors, including individual context elements, morphine dose, and locomotor effects of morphine, interact to affect morphine-context associations in both sexes. We find that while both male and female mice exhibit similar preference for morphine-paired contexts, males have slightly higher innate bias toward some of the contextual features of the testing apparatus. We further find that females display 1) greater dose-dependent morphine-induced hyperlocomotion than males and 2) greater locomotor sensitization than males when sufficient contextual information is provided for CPP to occur. Given the sex differences in the acute and repeated effects of morphine and withdrawal, we found that the locomotor effects of repeated morphine administration and withdrawal are sex-dependently correlated with the strength and stability of morphine-context associations. Specifically, acute morphine sensitivity predicted high CPP expression in females, while locomotor sensitization predicted lower CPP expression in males. Notably, we find that mice are likely in a state of morphine withdrawal during the morning saline conditioning (C2-C4) and P1 test sessions, as their locomotion is decreased. Withdrawal-induced hypolocomotion in males may have interfered with the expression of CPP, an effect that does not seem to appear in females. Together, these results suggest that different aspects of subjective experience of morphine intoxication and withdrawal are critical for morphine abuse-related behaviors in males and females.

In people with OUD, context is a major trigger for opioid use and relapse. Here we examined how contextual richness affected CPP, one of the major paradigms used to model OUD-related behavior. We found that there were innate sex-dependent biases in preference for contextual features, including wall pattern and especially floor texture (Fig. 1C**,D**), which is agreement with previous findings showing that tactile cues may be more effective for CPP conditioning compared to visual cues (Cunningham *et al*., 2006). Even so, neither compartment bias nor (randomly assigned) compartment that morphine was paired with affected basal locomotion or morphine-induced hyperlocomotion (Fig. 1E**,F**; Fig. 2). Males and females similarly acquired CPP in the high context assay (Fig. 4B), even though males had a higher initial preference for the textured floor compartment, suggesting that these innate preferences may not affect males’ and females’ ability to use contextual features to distinguish compartments necessary for making opioid-context associations, or our ability to measure this. This is interesting given some previous evidence that innate preference for certain features (such as wall color) can modulate CPP (Roma & Riley, 2005). Further, our results suggest that sufficient contextual information was present in the high context version, but not low context version, of our paradigm to elicit CPP in both sexes. This is in line with previous findings showing that factors including greater tactile information (Vezina & Stewart, 1987) and higher morphine concentration (until a sedative or aversive dose is reached) (Contarino *et al*., 1997) enhance morphine CPP in male rats. While CPP acquisition was similar in males and females for both the low and high context versions of our assay, our observations regarding innate biases toward context features highlight the importance of attending to possible sex differences in the experimental design of studies utilizing both sexes.

Using both the low and high context versions of the CPP assay allowed us to probe the importance of context and its strength in the locomotor effects of morphine, as well as the nature of the relationship between the locomotor effects of morphine with morphine-context associative learning and memory (both of which we found to depend on sex). We observed that males and females had similar dose-dependent locomotor responses to acute morphine administration across the wide range of doses used (Fig. 2), however, basal and morphine-induced locomotion were greater in females (as illustrated across **Figs. 1-3**). This is unsurprising given the evidence in both humans and rodents that morphine and other opioids can have diverging physiological effects depending on sex (Fullerton *et al*., 2018, 2022; Averitt *et al*., 2019). For example, women require more morphine than men to produce a similar degree of analgesiaClick or tap here to enter text., are more susceptible to overdose death from opioid prescriptionClick or tap here to enter text., and are more likely to escalate use of prescription opioids to the more potent and harmful illicit opioids, such as heroin (Cepeda & Carr, 2003; Ait-Daoud *et al*., 2019). In addition to the sex differences in pharmacological effects of acute morphine, contextual elements in the CPP assay contributed to an observed sex difference in locomotor sensitization to morphine.

Along with higher basal locomotion, females displayed steeper decreases in saline locomotion across sessions *and* steeper increases in morphine-induced hyperlocomotion across sessions in the low context assay (Fig. 3A**,C****,E**). These features changed in a dose-dependent manner in both sexes, leading to similar dose-dependent locomotor sensitization scores in females and males in the low context environment in which CPP was not achieved. In the high context assay, while morphine-induced hyperlocomotion was still greater in females than males, females displayed similar saline locomotion as males, leading to higher locomotor sensitization scores in females than males (Fig. 3B**,D****,F**). Therefore, this context-dependent sex difference in sensitization seems to be particularly driven by an effect of context on females’ locomotion during saline sessions. These data implicate sex differences in the underlying organization and function of brain circuits encoding and relaying context information. For example, dopamine 1 receptor (D1R)-expressing and dopamine 2 receptor (D2R)-expressing neurons in the nucleus accumbens shell (NAcSh) have been shown to play distinct roles on the induction and later escalation of locomotor sensitization to psychostimulants (Kai *et al*., 2015). It is possible that sexually dimorphic expression or modulation by dopamine or mu opioid receptors (Loyd *et al*., 2008), or sex-dependent innervation from context-encoding regions such as the hippocampus, affect the recruitment of these NAcSh neuronal subpopulations to contribute to context-dependent sex differences.

Studies have classically used saline session locomotion as a normalization factor to evaluate morphine locomotor sensitivity (inferring that locomotor sensitization is driven by morphine-induced hyperlocomotion), however here we find in reporting the raw saline locomotion that mice may be in a state of withdrawal during these sessions in a manner that affects locomotor sensitization and CPP. We found that mice did not sensitize to repeated morphine administration at a dose of 5 mg/kg, but they did in a dose-dependent manner across 10, 20, and 50 mg/kg doses (Fig. 3C**,D**). In addition, there was no or little decrease in locomotion across daily saline sessions for the 5 mg/kg morphine dose, but saline locomotion decreased across conditioning days to a steeper extent at the higher doses in a dose-dependent manner (Fig. 3A**,B**) to a similar degree as the morphine-induced increase. These results suggest that no or little locomotor habituation occurred in the saline-paired compartment of the CPP apparatus and that saline hypolocomotion was related to morphine hyperlocomotion. As such, hypolocomotion during saline sessions on C2-C4 in mice at ≥10 mg/kg morphine indicates that the mice were in a state of increased withdrawal across daily saline sessions. This interpretation is further supported by a dose-dependent hypolocomotive phenotype on the CPP test the day after the last morphine conditioning day (P1) that recovered 10 days later on P11 (Fig. 5; Fig. 6D**,E**). The potential for withdrawal being paired with a specific context may increase the salience of the saline-paired context and promote discrimination between contexts required for a place preference for the morphine-paired compartment of the CPP apparatus. A recent study supports the notion that the saline (withdrawal)-paired context is critical for CPP, as it showed that place cells in the hippocampus encoding the morphine-paired compartment during a CPP paradigm show stable activity across conditioning sessions and post-tests, while the cells encoding the saline-paired compartment diminish their activity and place cell function across conditioning; thus the discrimination between the two contexts was characterized primarily by neuroplastic changes during the saline conditioning sessions (Sun & Giocomo, 2022).

Our results further highlight that the reinforcing properties of a drug should not be inferred solely from its locomotor sensitivity scores. For example, both males’ and females’ locomotion suggest that the animals were more sensitive to 20 than 10 mg/kg morphine, while CPP scores were similar between the doses. However, we found that relationships between the locomotor effects of morphine intoxication/withdrawal and CPP were opposing between males and females (Fig. 6G**,H**; Fig. 7A). Morphine locomotor sensitization across conditioning and locomotion during withdrawal on P1 (but not locomotion on P11 after the withdrawal-related hypolocomotor phenotype recovered) were negatively associated with males’ CPP scores (Figs. 4, 6, 7), suggesting that a state of withdrawal impaired the acquisition or expression of CPP in males. The enhanced sensitization score in high context females compared to males was driven by females’ increase in morphine locomotion rather than decrease in saline locomotion, suggesting a specific role for the positive locomotor effect of morphine in females (Fig. 3). Indeed, the positive acute and sensitizing locomotor effects of morphine were correlated with a higher CPP score in females in the high context assay (Fig. 4C). Notably, these relationships had similar directionality for the low context females, even though the groups on average did not show a robust CPP, implying that some mice and/or some degree of CPP may have developed in a subset of females even when minimal context features were present (Fig. 7).

Altogether, these findings have implications for sex differences in the organization and function of circuits and neurotransmitter/neuromodulatory systems that play a role in locomotion, reward, and aversion, as these are involved during states of morphine intoxication and withdrawal. Furthermore, there may be sex differences in the ways that environmental features influence the interoceptive effects of drugs and the strength and stability of related context associations. While the roles of factors influencing drug-context learning and memory have been assessed in male rodents, this has been understudied in females. However, the ample sex differences in the organization and function of the hippocampus and nucleus accumbens (Nestler, 2001; Forlano & Woolley, 2010; Kutlu & Gould, 2016; Yagi *et al*., 2022) increase the likelihood of sex differences in how environmental features affect the interoceptive effects of drugs, as well as the strength and stability of context-dependent drug associations (Forlano & Woolley, 2010; Kobrin *et al*., 2016; Kutlu & Gould, 2016; Alvandi *et al*., 2017; Nazari-Serenjeh *et al*., 2021; Trott *et al*., 2022). For example, females have greater distal dendritic spine density and proportion of large spines in the NAc compared to males, which may enhance the strength or efficiency of glutamatergic synapses from the hippocampus and other regions (Forlano & Woolley, 2010). Further, several studies suggest that there are sex differences in memory for aversive stimuli and neural activation in response to the recall of those memories. Male and female rats show distinct patterns of functional connectivity in limbic regions involved in fear memory, with female rats displaying greater activation in the frontal cortex and dorsal CA1 region of the hippocampus (Yagi *et al*., 2022) even though male rats may be more sensitive to contextual fear in general (Trott et al. 2022). These anatomical and functional sex differences that affect acquisition, encoding, and expression of the relationships between context and aversive outcomes may also contribute to drug-context associations.

## 5 Conclusions

We found that females and males displayed similar dose-dependent locomotor responses to acute and repeated morphine exposure, as well as morphine-context associations. However, there were intriguing sex differences in the effects of context on locomotor sensitivity to morphine, as well as in the contributions of morphine intoxication and withdrawal phases of conditioning on locomotor sensitization and the relationships between the locomotor effects of morphine and the associations formed between the drug and the context in which it was experienced. These results have implications for understanding observed sex differences in humans in responses to acute and chronic exposure to morphine and other opioid drugs. They also may better inform sex-specific strategies for pharmacological treatment approaches to curb opioid seeking and taking behaviors, including relapse, which are highly influenced by context and withdrawal severity.

